# Sex chromosome pairing mediated by euchromatic homology in *Drosophila* male meiosis

**DOI:** 10.1101/854877

**Authors:** Christopher A. Hylton, Katie Hansen, Andrew Bourgeois, John E. Tomkiel

## Abstract

To maintain proper ploidy, haploid sex cells must undergo two subsequent meiotic divisions. During meiosis I, homologs pair and remain conjoined until segregation at anaphase. *Drosophila melanogaster* spermatocytes are unique in that the canonical events of meiosis I including synaptonemal complex (SC) formation, double-strand DNA breaks, and chiasmata are absent. Sex chromosomes pair at intergenic spacer sequences within the heterochromatic rDNA while euchromatin is required to pair and segregate autosomal homologies, suggesting that pairing may be limited to specific sequences. However, previous work generated from genetic segregation assays or observations of late prophase I/prometaphase I chromosome associations fail to differentiate pairing from conjunction. Here, we separately examined the capability of X euchromatin to pair and conjoin using an rDNA-deficient X and a series of *Dp(1;Y)* chromosomes. Genetic assays showed that duplicated X euchromatin can substitute for endogenous rDNA pairing sites. Segregation was not proportional to homology length, and pairing could be mapped to nonoverlapping sequences within a single *Dp(1;Y)*. Using fluorescent *in situ* hybridization (FISH) to early prophase I spermatocytes, we showed that pairing occurred with high fidelity at all homologies tested. Pairing was unaffected by the presence of X rDNA, nor could it be explained by rDNA magnification. By comparing genetic and cytological data, we determined that centromere proximal pairings were best at segregation. Segregation was dependent on the conjunction protein Stromalin in Meiosis while the autosomal-specific Teflon was dispensable. Overall, our results suggest that pairing may occur at all homologies, but there may be sequence or positional requirements for conjunction.

**ARTICLE SUMMARY:** *Drosophila* males have evolved a unique system of chromosome segregation in meiosis that lacks recombination. Chromosomes pair at selected sequences suggesting that early steps of meiosis may also differ in this organism. Using Y chromosomes carrying portions of X material, we show that pairing between sex chromosomes can be mediated by sequences other than the previously identified rDNA pairing sites. We propose that pairing may simply be homology-based and may not differ from canonical meiosis observed in females. The main difference in males may be that conjunctive mechanisms that join homologs in the absence of crossovers.

## INTRODUCTION

Meiosis is the highly conserved process comprised of two cell divisions that produce four haploid daughter cells from a single diploid parent cell. To ensure an equal distribution of homologous chromosomes to gametes, homologs must locate each other, pair, conjoin, and segregate with high fidelity. Several events have been identified that aid in homolog pairing, but the mechanisms of partner recognition remain enigmatic. Multiple plant species create a chromosome “bouquet” by clustering and imbedding all telomeres into the inner nuclear membrane thereby confining homolog identification and pairing to a smaller region of the nucleus (Bahler *et al*. 1993). *Caenorhabditis elegans* uses microtubule/dynein-mediated movements through linkages to telomeric chromosomal sites deemed “pairing centers” which are thought to facilitate interactions between homologs (MacQueen *et al*. 2005; Sato *et al*. 2009; Wynne *et al*. 2012). The budding yeast *Saccharomyces cerevisiae* establishes DNA sequence-independent associations between homologous centromeres prior to bouquet formation to enhance the odds that homologous pairs of kinetochores attach to the correct spindle pole (Kemp *et al*. 2004). Despite progress in understanding the mechanisms that aid in homolog association, the molecular basis of pairing itself remains poorly understood.

Recombination appears to play an essential role in pairing in some systems. During meiosis I of *S. cerevisiae*, the formation of double-stranded breaks, a prerequisite for recombination, occurs prior to homolog synapse initiation. In *spo11* yeast that lack double strand breaks, homologs fail to synapse (Giroux, Dresser, and Tiano 1989; Weiner and Kleckner 1994), which indicates that the homology search achieved by single-stranded DNA during recombination in yeast is required for homolog pairing and synapsis. In contrast, *mei-W68* and *mei-P22 Drosophila* females that lack double-strand breaks and crossing over assemble SC indicating recombination is not required for pairing and synapsis (McKim *et al*. 1998). Taken together, these results reveal that while some species require recombination for pairing, other species have evolved separate recombination-independent mechanisms to pair and segregate homologs.

Male *Drosophila*, which completely lack recombination, have two genetically separable pathways to pair and segregate chromosomes. One pathway is specific for the sex chromosomes and the other for the autosomes. Sex chromosomes pair at specific sites, originally termed collochores, that were identified based on the observation that certain regions of the X and Y remain associated at prometaphase I and metaphase I (Cooper 1959). Potential pairing sites were identified in the repetitive heterochromatic region near the centromere of the X chromosome and near the base of the short arm of the Y chromosome. These two regions contain sequence homology of the rDNA genes, which contain 200-250 tandem copies of the genes for the ribosomal subunits (Ritossa 1976). Males with rDNA-deficient X chromosomes exhibit high levels of X-Y nondisjunction (NDJ). A transgenic copy of the rDNA gene on the X restores disjunction (McKee and Karpen 1990). The 240 bp intergenic spacer (IGS) region located upstream of each 18S and 28S rDNA repeat is necessary and sufficient for pairing (McKee, Habera, and Vrana 1992). To date, the IGS sequences are the only pairing sites identified in any organism.

In contrast to the sex chromosomes which lack euchromatin, autosomes pair at sequences that are distributed throughout the euchromatin, and both the amount and chromosomal location of euchromatic homology may be important for conjunction (McKee, Lumsden, and Das 1993). Cytological and genetic tests show that autosomes with only heterochromatic homology fail to segregate from each other at meiosis I (Yamamoto 1979; Hilliker, Holm, and Appels 1982). These studies suggested that autosomal heterochromatin lacked pairing ability.

Because these conclusions were largely derived from observations of chromosome associations during late prophase I to prometaphase I, sequences were only defined as pairing sites if they had the ability to remain conjoined. The initial interactions needed for homolog recognition and pairing occur premeiotically, however, and at these later stages, many interactions may have already been resolved. Thus, the previously defined “pairing sites” may really represent regions that remain conjoined and may not necessarily represent all sequences involved in pairing.

Direct observations of pairing provide a more accurate assessment of pairing sites. Meiotic pairing is separable from homolog associations that occur in somatic cell (“somatic pairing”). Homologs are not paired at the earliest stage that germline cells can be distinguished in the embryo, but then begin to associate in gonial cells prior to meiosis (Joyce *et al*. 2013). Examination of early prophase I pairing in vivo using the GFP-Lac repressor/lac operator system, found that homologs were paired at each of 13 different single autosomal loci (Vazquez, Belmont, and Sedat 2002). In agreement with earlier studies, this shows that many autosomal sequences can pair. Heterochromatic homologies also pair with similar kinetics, as shown by *in situ* hybridizations to autosomal satellite repeats (Tsai, Yan, and McKee 2011).

Distinct from pairing, conjunction refers to the ability of paired homologs to remain coupled during prophase I condensation and prometaphase/metaphase I spindle-mediated movements. Teflon (Tef), Modifier of Mdg in Meiosis (MNM), and Stromalin in Meiosis (SNM) have all been shown to be required for conjunction of the autosomes, while sex chromosome conjunction requires only MNM and SNM (Tomkiel, Wakimoto, and Briscoe 2001; Thomas *et al*. 2005). MNM and SNM localize to the rDNA on the sex chromosomes (Thomas *et al*. 2005) specifically at the rDNA IGS (Thomas and McKee 2007). Potential MNM/SNM/Tef and MNM/SNM complexes may regulate autosomal and sex chromosome conjunction, respectively, holding paired homologs together until anaphase I (Thomas *et al*. 2005; Thomas and McKee 2007). Recently, super resolution microscopy and temporally expressed transgenes showed that MNM and SNM are required to maintain conjunction but cannot establish pairing themselves (Sun *et al*. 2019). Thus, while the 240 IGS pairing sites on the X and Y certainly have the ability to mediate pairing and may serve as a site for conjunction protein binding, they may not be the only sequences with the ability to pair. It remains to be examined if other sequence homologies can pair but lack the ability to stabilize conjunction.

Here, we directly examine pairing and its relationship to conjunction. We describe a system to examine sex chromosome pairing during early prophase I at homologies other than the IGS repeats. We show that X euchromatic sequences placed on the Y chromosome are able to pair and in some cases facilitate conjunction and segregation of sex chromosomes in the absence of X chromosome rDNA. This system allowed us to identify sequences capable of pairing, to ask how much homology is sufficient for pairing, and to determine whether the location of homology is important for pairing and conjunction.

## MATERIALS AND METHODS

### *Drosophila* Stocks and Crosses

*Drosophila* were raised on a standard diet consisting of cornmeal, molasses, agar, and yeast at 23°C. *Dp(1;Y)* chromosomes (Cook *et al*. 2010) and *Df(tef)803Δ15* (Arya *et al*. 2006) are previously described. The *tef ^z3455^*, *snm^z0317^*, *snm^z2138^*, *mnm^z5578^*, *mnm^z3298^*, and *mnm^z3401^* alleles were originally obtained from the C. Zuker laboratory at the University of California at San Diego (Wakimoto, Lindsley, and Herrera 2004) and are previously described (Tomkiel, Wakimoto, and Briscoe 2001; Thomas *et al*. 2005). All other stocks were obtained from the Bloomington Stock Center (Gramates *et al*. 2017).

### Genetic Assays of Meiotic Chromosome Segregation

*In(1)sc^4L^sc^8R^* and *Df(1)X-1* are X chromosomes that have been reported to be rDNA-deficient. We found that *Df(1)X-1* X resulted in sterility in combination with the *Dp(1;Y)* Y chromosomes tested, and therefore used the *In(1)sc^4L^sc^8R^* X was selected for crosses. Segregation of *In(1)sc^4L^sc^8R^* from a *Dp(1;Y)* chromosome was monitored by crossing *In(1)sc^4L^sc^8R^ y^1^* / *Dp(1;Y)B^S^ Y y^+^* males to *y w sn; C(4)RM ci ey / 0* females. Offspring are scored as either normal (B^S^ y^+^ sn sons or y^1^ daughters), sex chromosome diplo-(B^S^ y^+^ females), or sex chromosome nullo-exceptions (y w sn males). The midpoint of the duplicated X euchromatin on each *Dp(1;Y)* was calculated by taking the average of the distal- and proximal-most estimations of breakpoints (Cook *et al*. 2010).

Fourth chromosome missegregation was monitored by the recovery of ci ey *nullo-4* progeny. In crosses involving *tef* mutations, males were made homozygous for the fourth chromosome mutation *spa* to allow monitoring of both *nullo-4* and *diplo-4* progeny.

### Probe Design

Probe pools were generated to selected sequences at a density of 10 probes/Kb and a complexity of ∼10,000 probes per pool (Arbor Biosciences, Ann Arbor, MI). Triple-labeled Atto-594 oligonucleotide probes were generated to sequences present on both *In(1)sc^4L^sc^8R^* and the following *Dp(1;Y)* chromosomes:

*Dp(1;Y)BSC76* – X salivary gland chromosome bands 2E1-3E4, bp 2606837-3606837
*Dp(1;Y)BSC185* – X salivary gland chromosome bands 12A4-12F4, bp 3824004 - 14826069
*Dp(1;Y)BSC11* – X salivary gland chromosome bands 16F7-18A7, bp 18193946-19193592.

A triple-labeled Atto-488 probe was generated to bp 20368577-21368577 (56F-57F) on chromosome 2. An Atto-488 probe (Eurofins MWG Operon, Louisville, KY) was synthesized to the Y-specific AATAC heterochromatic repeat (Lohe and Brutlag 1987).

## FISH

Slides of testis tissue were processed for FISH using a modification of the protocol as described (Beliveau, Apostolopoulos, and Wu 2014). Testes from larvae (Pairing Assay) or pharate adults (NDJ Assay) were dissected in Schneider’s *Drosophila* media (GIBCO BRL, Gaithersburg, MD). Tissue was transferred to a drop of Schneider’s on a silanized coverslip and gently squashed onto a Poly-L-Lysine coated slide (Electron Microscopy Sciences, Hatfield, PA).

Coverslips were immediately removed after freezing in liquid nitrogen. Tissue was fixed in 55% methanol/25% acetic acid for 10 min followed by 10 min dehydration in 95% ethanol. Slides were processed immediately or stored for up to 1 week at 4°C.

For hybridizations, slides were rehydrated in 2X saline-sodium citrate/Tween-20 (SSCT) at room temperature for 10 min. Membranes were permeabilized and DNA denatured by incubation in 50% formamide/2X SSCT for 2.5 min at 92°C then 60°C for 20 min. Slides were rinsed in 1X phosphate buffered saline (PBS) for 2 min and allowed to dry. 5 µl of probe master mix containing 12.5 µl hybrid cocktail (50% dextran sulfate, 20X SSCT), 12.5 µl formamide, 1 µl of 10 mg/ml RNase, 2 µl of probe 1 (5 pmol/µl), and 2 µl of probe 2 (5 pmol/µl) was pipetted directly onto a silanized 18 x 18 mm coverslip, which was placed on the tissue and sealed with rubber cement. Slides were heated at 92°C for 2 min to denature the DNA then incubated in a damp chamber at 42°C for >18 hours. Following incubation, coverslips were removed, and slides were incubated in 2X SSCT at 60°C for 20 min, 2X SSCT at RT for 10 min, and 0.2X saline-sodium citrate (SSC) at RT for 10 min to remove unbound probe. DNA was stained with 1 µg/µl 4’,6-diamidino-2-phenylindole (DAPI) (Sigma, St. Louis, MO) and tissues mounted in ProLong Gold antifade (Invitrogen, Carlsbad, CA). Probes were visualized using a Keyence BZ-X700 Fluorescence Microscope. S1-S2 spermatocytes were selected based on size (10 to 20 µm), and signals were scored as paired when within 0.8 µm (Beliveau, Apostolopoulos, and Wu 2014).

### Estimation of the ability of paired sequences to direct segregation

To determine how frequently pairing led to disjunction, we assumed that chromosomes that did not pair would segregate at random. First, we determined the pairing frequency from FISH assessment of S1-S2 cells (= % Paired). We then cytologically determined the frequency of secondary spermatocytes and spermatids in which the X and Y had segregated to opposite poles at meiosis I (= % NDJ). We assumed that this latter frequency represented meiocytes in which XY pairings underwent normal segregation, plus half the frequency of random disjunctions that resulted when the X and Y failed to pair. Based on this assumption, we calculated the percent of cells in which pairing of XY chromosomes led to normal disjunction as: Paired then disjoined = (% Paired – [%NDJ – (1/2 % Unpaired)]) / % Paired.

### rDNA Magnification Assay

rDNA magnification was assessed by crossing *In(1)sc^4L^sc^8R^ y^1^* / *Y* males (Cross A) or *In(1)sc^4L^sc^8R^ y^1^* / *Dp(1;Y)B^S^ Y y^+^ BSC76* males (Cross B) to *C(1)RM*, *y w f* / *y^+^ Y* females. Fifty *In(1)sc^4L^sc^8R^ y^1^* / *y^+^ Y* sons generated from cross A or B were then crossed to y w sn females to determine sex chromosome NDJ. NDJ was calculated amongst progeny of each father, and distributions of NDJ frequencies were compared by one-way ANOVA.

### Data Availability

All strains are available on request. The authors affirm that all data necessary for confirming the conclusions of the article are present within the article, figures, and tables.

## RESULTS

### Euchromatic homology directs segregation of the X from the Y

We developed a system to ask if euchromatic homologies could direct pairing and segregation of the sex chromosomes utilizing a series of *Dp(1;Y)* chromosomes (Cook *et al*. 2010) and the rDNA-deficient *In(1)sc^4L^sc^8R^* X chromosome that is missing the sex chromosome pairing sites. Each *Dp(1;Y)* chromosome contains a unique segment of X euchromatin. The size and position of the duplicated homology with the X chromosome partner also varies (Figure 1). We reasoned if the euchromatic homology was sufficient to pair, conjoin, and direct segregation of the sex chromosomes, then *In(1)sc^4L^sc^8R^ / Dp(1;Y)* males would produce fewer exceptional progeny than *In(1)sc^4L^sc^8R^ / Y* males.

**Figure 1.**
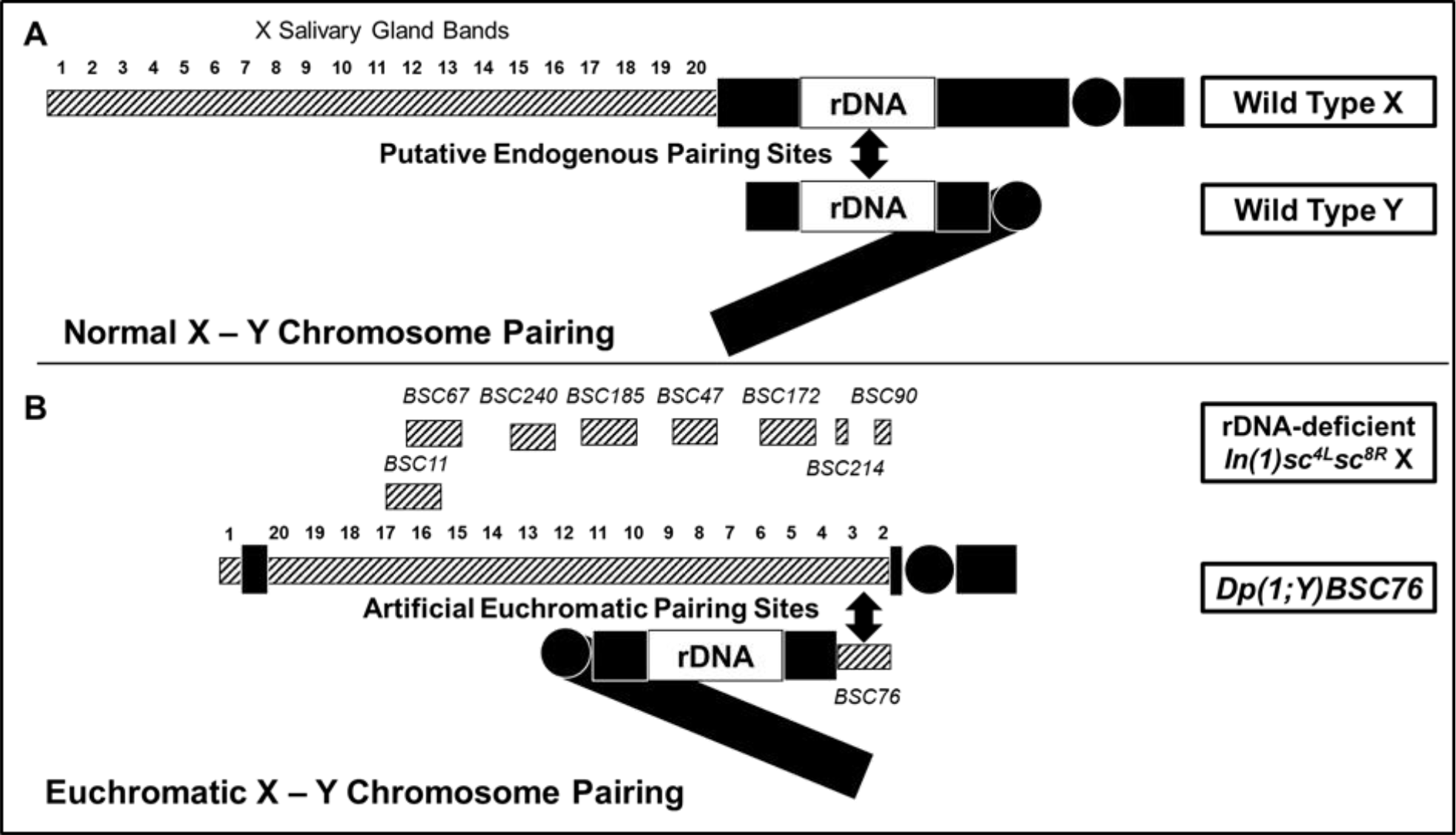
Normal X-Y pairing vs. pairing at euchromatin (Hatched Boxes). (A) Wildtype showing rDNA pairing sites. (B) *In(1)sc^4L^sc^8R^* X lacking rDNA. The locations of the X duplications on the collection of *Dp(1;Y)s* tested are indicated above the X. *Dp(1;Y)BSC76* is shown paired with its euchromatic homology on the X.

As a metric of segregation, we monitored NDJ of the sex chromosomes among progeny of *In(1)sc^4L^sc^8R^* / *Dp(1;Y)* males. Direct comparisons of the behaviors of the different *Dp(1;Y)* males are complicated as the viabilities of *Dp(1;Y)*-bearing sons differ greatly (data not shown), most likely a result of gene dosage imbalance contributed by the X duplications. To directly compare the behaviors of different *Dp(1;Y)* chromosomes, we considered only two classes of progeny that were genetically identical from all crosses. X/0 sons were used as a metric of sex chromosome NDJ, and X/X daughters were used as a metric of normal disjunction. We used the ratio of (X / 0) / (X / X + X / 0) as an estimate for the frequency of missegregation of sex chromosomes in each class of test males, and for the remainder of the manuscript, sex chromosome NDJ will be determined as such. We found that some of the *Dp(1;Y)*s were better at segregating from the X chromosome (Table 1). The ability to segregate was not related to the length of the duplicated X euchromatin sequence (Figure 2A), however we noted a relationship between proper X-Y segregation and the chromosomal location of X homology. When the homologous sequences on the inverted X chromosome were closer to the centromere less NDJ was observed (Figure 2B). As a control, chromosome 4 segregation was also monitored to determine if X duplicated material itself generally perturbed chromosome segregation due to effects of aneuploidy. 4^th^ chromosome NDJ was less than 1% in each of the *Dp(1;Y)*-bearing males tested, indicating that none of the *Dp(1;Y)*s increased autosomal NDJ (data not shown).

**Figure 2.**
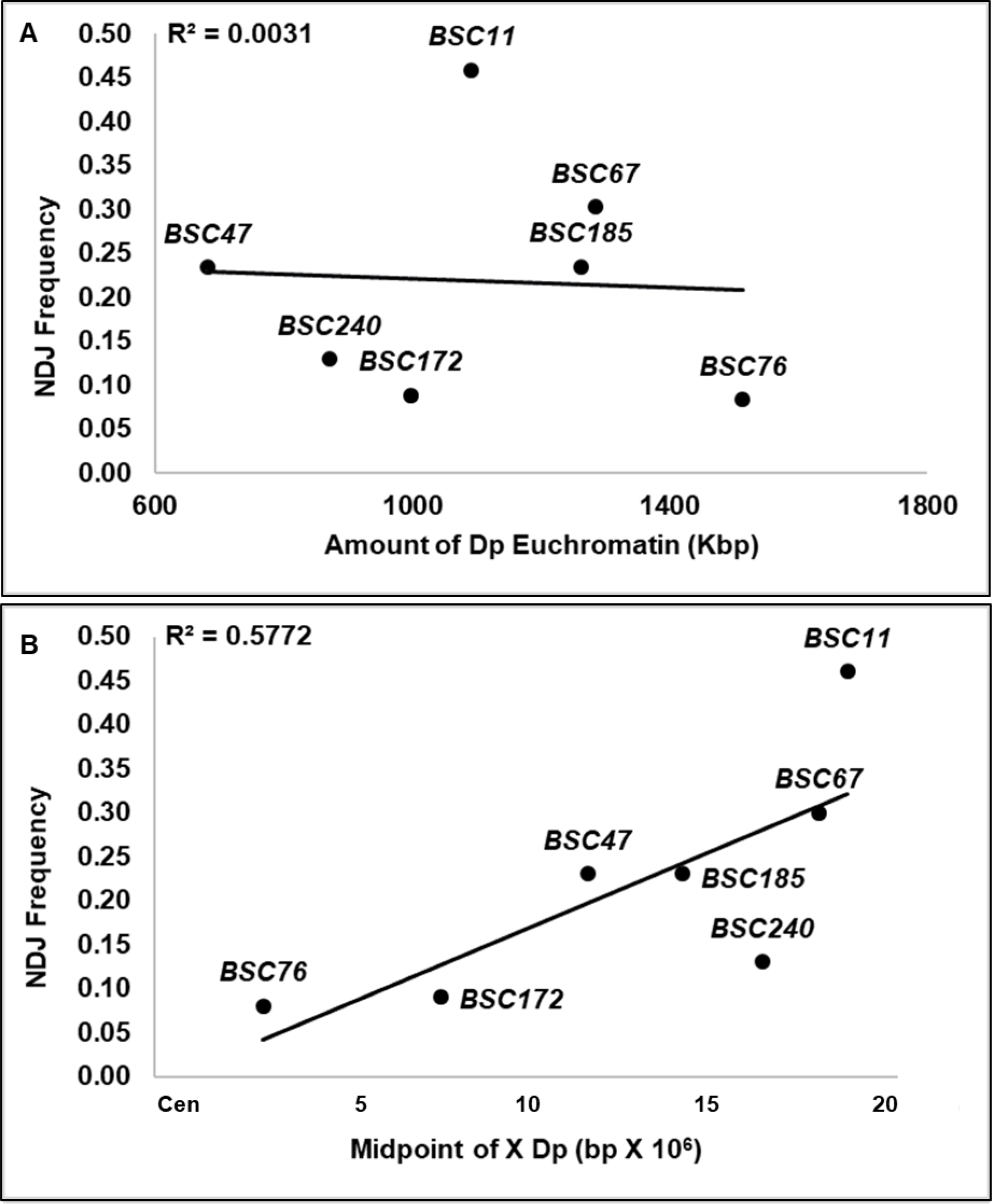
Sex chromosome NDJ frequencies among progeny of *In(1)sc^4L^sc^8R^ / Dp(1;Y)BSC* males versus (A) euchromatic homology length and (B) genomic sequence position of the X homology.

**TABLE 1.** Frequency of XY NDJ among progeny from In(1)sc^4L^sc^8R^ / Dp(1;Y) males

| Sperm genotype: |  | X | *Dp(1;Y)* | *X/Dp(1;Y)* | 0 | 0/(X+0) |
| --- | --- | --- | --- | --- | --- | --- |
| Paternal Y | X region duplicated on Y * |  |  |  |  |  |
| *y+Y* | - | 925 | 434 | 52 | 579 | 0.38 |
| *Dp(1;Y)BSC76* | 2E1-3E4 | 321 | 51 | 0 | 29 | 0.08 |
| *Dp(1;Y)BSC172* | 7A3-7D18 | 319 | 41 | 0 | 31 | 0.09 |
| *Dp(1;Y)BSC47* | 10B3-11A1 | 387 | 35 | 2 | 118 | 0.23 |
| *Dp(1;Y)BSC185* | 12A4-12F4 | 421 | 75 | 2 | 129 | 0.23 |
| *Dp(1;Y)BSC240* | 14A1-15A8 | 660 | 59 | 1 | 98 | 0.13 |
| *Dp(1;Y)BSC67* | 15F4-17C3 | 307 | 35 | 1 | 133 | 0.30 |
| *Dp(1;Y)BSC11* | 16F7-18A7 | 495 | 69 | 19 | 419 | 0.46 |

\* Salivary gland chromosome bands

We conclude that the duplicated X euchromatin on the Y chromosome is capable of facilitating pairing, conjunction, and segregation of the sex chromosomes, and that the ability to do so is related to underlying sequences and/or chromosomal position. However, a potential caveat to our interpretation is that our genetic metric may be influenced by ‘meiotic drive’, a phenomenon that results in the unequal recovery of reciprocal meiotic products. Meiotic drive is induced by a failure of sex chromosome pairing in male flies, and drive strength is directly proportional to the pairing frequency (McKee 1984). Although termed ‘meiotic drive’, this process has been shown to result in a post-meiotic differential elimination of sperm dependent on chromatin content (Peacock, Miklos, and Goodchild 1975). Thus, it was a formal possibility that the differences we had observed could somehow result from differential effects of the various *Dp(1;Y)* chromosomes on meiotic drive. To avoid this potential complication, we turned to a direct cytological assessment of chromosome behavior in meiosis.

We used FISH with X- and Y-specific probes to directly assess the outcomes of meiosis in secondary spermatocytes and onion stage spermatids. An Atto-594 (Red) X chromosome probe labels an X euchromatic sequence, while an Atto-488 (Green) Y chromosome labels the unique AATAC heterochromatic repeat. Segregation frequencies of the sex chromosomes were determined by examining related pairs of secondary spermatocytes, or related tetrads of spermatids (Table 2). This analysis confirmed our conclusions based on our genetic observations, that the fidelity of segregation from *In(1)sc^4L^sc^8R^* varied among tested *Dp(1;Y)*s, and this variation was related to proximity of the homology to the X centromere (Figure 3).

**Figure 3.**
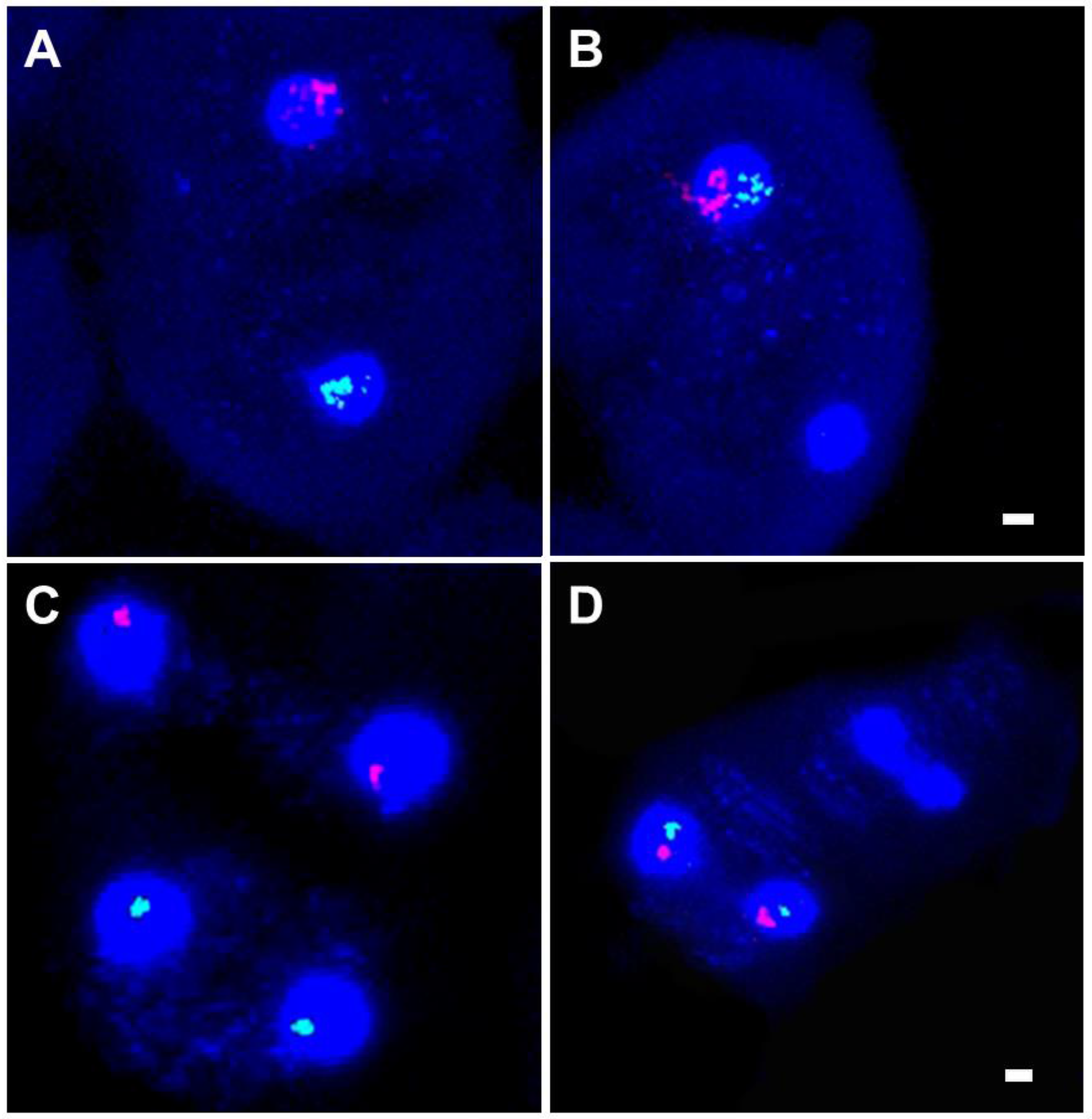
FISH examination of *In(1)sc^4L^sc^8R^* / *Dp(1;Y)BSC76* disjunction in DAPI-stained spermatocytes using an X probe (Red) and an AATAC repeat Y probe (Green). (A) Normal XY segregation during meiosis I and (B) meiosis I NDJ. (C) Meiosis II division after a normal meiosis I division and (D) after a meiosis I NDJ. Scale bar = 2 µm.

**TABLE 2.** XY NDJ frequencies as determined by FISH.

| X | Y | # Meioses Scored | XY NDJ |
| --- | --- | --- | --- |
| Canton S | Canton S | 206 | 0.00 |
| *In(1)sc<sup>4L</sup>sc<sup>8R</sup>* | *y+* Y | 200 | 0.33 |
| *In(1)sc<sup>4L</sup>sc<sup>8R</sup>* | *Dp(1;Y)BSC76* | 307 | 0.11 |
| *In(1)sc<sup>4L</sup>sc<sup>8R</sup>* | *Dp(1;Y)BSC185* | 214 | 0.22 |
| *In(1)sc<sup>4L</sup>sc<sup>8R</sup>* | *Dp(1;Y)BSC11* | 237 | 0.30 |
| *In(1)sc<sup>4L</sup>sc<sup>8R</sup>* | *Dp(1;Y)BSC90* | 201 | 0.12 |
| *In(1)sc<sup>4L</sup>sc<sup>8R</sup>* | *Dp(1;Y)BSC214* | 206 | 0.08 |

While these observations clearly suggest that the various *Dp(1;Y)* chromosomes were pairing with the rDNA-deficient X, they do not address where this pairing might be occurring. It is known that in the presence of structurally altered Y chromosomes, a process termed rDNA magnification can be induced (Tartof 1974). This process involves stable increases and/or decreases in rDNA copy number on an rDNA-deficient X via unequal sister chromatid exchange (Ritossa 1968). Although the *In(1)sc^4L^sc^8R^* chromosome is reportedly deleted for all of the rDNA, one or more cryptic rDNA cistrons could be potentially induced to magnify and restore XY pairing via the endogenous rDNA pairing sites. As few as six copies of the rDNA intergenic spacer repeats may restore pairing between the X and the Y (Ren *et al*. 1997), thus it was important to determine if our results could be explained by rDNA magnification rather than pairing outside the rDNA. To test for rDNA magnification, we provided potential magnification conditions by passing an *In(1)sc^4L^sc^8R^* X through a male bearing a *Dp(1;Y)*. We chose the *Dp(1;Y)* that exhibited the highest fidelity of segregation, *Dp(1;Y)BSC76*, as this would be predicted to show the greatest amount of magnification, if it were indeed occurring. We recovered the potentially amplified X chromosomes in sons, and genetically tested their ability to segregate from the Y. As a control, we tested genetically identical males which had received an *In(1)sc^4L^sc^8R^* that had not been exposed to potentially magnifying conditions. If magnification was occurring, then we expected that sons bearing the potentially magnified *In(1)sc^4L^sc^8R^* would demonstrate improved segregation of the sex chromosomes relative to the controls. For each test, we scored progeny of 50 males. No statistical difference was found between the two classes (ANOVA, F value = 1.76527; *p* value = 0.17475) (Figure 4, Table 3). We conclude that the ability of a *Dp(1;Y)* to segregate from an rDNA-deficient *In(1)sc^4L^sc^8R^* is not a consequence of rDNA magnification and likely reflects pairing between X euchromatic homologies.

**Figure 4.**
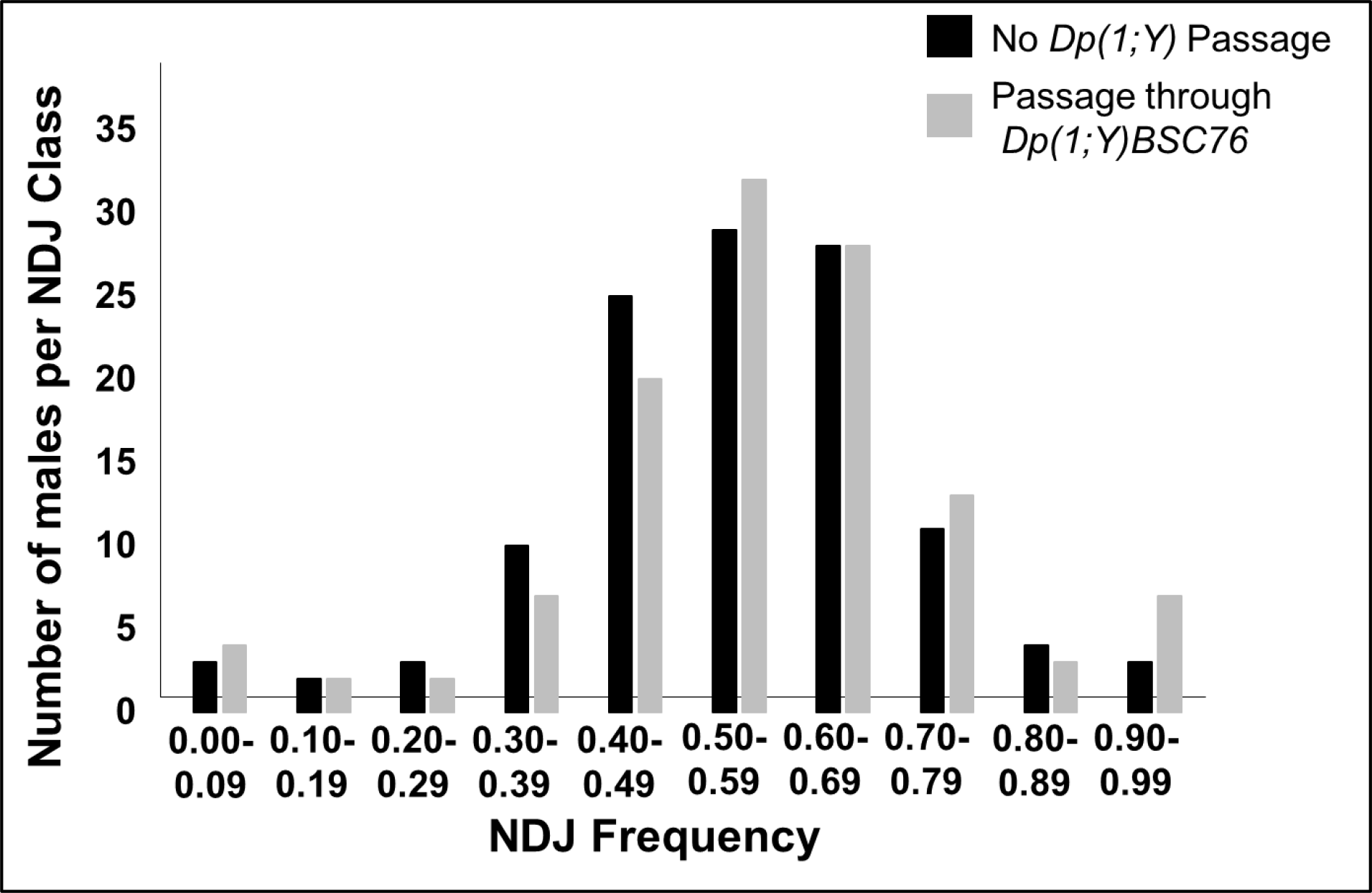
Test for rDNA magnification of *In(1)sc^4L^sc^8R^* in *Dp(1;Y)BSC76* males. Distributions of NDJ frequencies in sons of *In(1)sc^4L^sc^8R^* / *Dp(1;Y)BSC76* or *In(1)sc^4L^sc^8R^* / *Y* males.

**Figure 5.**
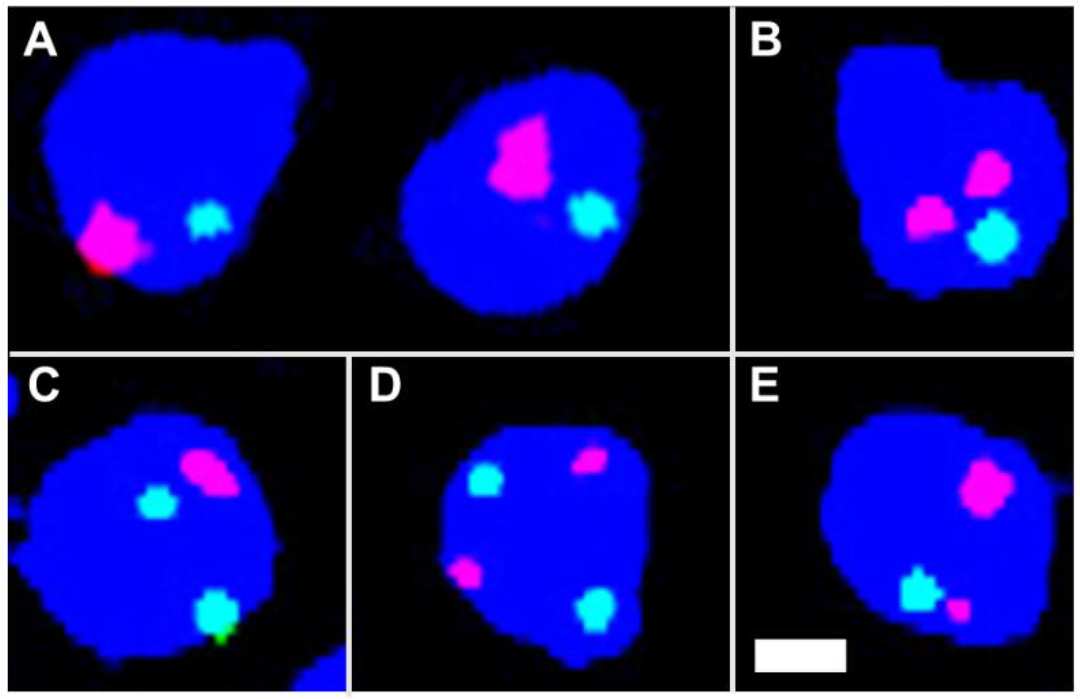
FISH examination of pairing in DAPI-stained S1-S2 primary spermatocytes using an X probe (Red) and chromosome 2 probe (Green). (A) Paired XY and paired chromosome 2 bivalents. (B) Unpaired XY and a paired chromosome 2 bivalent. (C) A paired XY bivalent and unpaired chromosome 2. (D) Both unpaired. (E) Sister chromatid separation from a paired XY bivalent. Scale bar = 2 µm.

**TABLE 3.** Frequency of XY NDJ among progeny from *In(1)sc^4L^sc^8R^ /* Y males after potential rDNA magnification.

| Sperm genotype: | X | Y | X/Y | 0 | (X/Y+0)/Total |
| --- | --- | --- | --- | --- | --- |
| No Magnification | 1962 | 747 | 86 | 2146 | 0.45 |
| Potential Magnification | 1668 | 563 | 74 | 2009 | 0.48 |

Homologies from various non-overlapping regions of the X chromosome enhanced segregation demonstrating that multiple sequences are capable of acting as pairing sites. Because no relationship between the length of the *Dp(1;Y)* and the ability to direct segregation was observed, we wanted to determine if these pairing site sequences were distributed randomly throughout the X euchromatin. To ask if we could potentially map a pairing site within a duplicated region, *Dp(1;Y)*s nested within the *Dp(1;Y)BSC76* euchromatic duplication were tested. The two smallest nonoverlaping duplications *Dp(1;Y)BSC90* and *Dp(1;Y)BSC214* were equally proficient at directing X-Y segregation albeit at a lower frequency than *Dp(1;Y)BSC76* (Table 4). These data suggest at least two different euchromatic segments within this one region are capable of pairing and directing X-Y segregation.

**TABLE 4.** Mapping segregational ability within Dp(1;Y)BSC76

|  |  |  | Sperm genotype: |  |  |  |  |
| --- | --- | --- | --- | --- | --- | --- | --- |
|  |  |  | X | *Dp(1;Y)* | *X/Dp(1;Y)* | 0 | 0/(X+0) |
| Paternal Y | X region duplicated on Y * | Homology ** (Kbp) |  |  |  |  |  |
| *Dp(1;Y)BSC76* | (2E1-2E2)-3E4 | 1514 | 1347 | 196 | 7 | 154 | 0.10 |
| *Dp(1;Y)BSC80* | (3A6-B1)-3E4 | 1100 | 1286 | 95 | 1 | 226 | 0.15 |
| *Dp(1;Y)BSC83* | (3B3-B4)-3E4 | 975 | 2105 | 420 | 14 | 280 | 0.12 |
| *Dp(1;Y)BSC84* | (3C2-C3)-3E4 | 803 | 1364 | 241 | 7 | 209 | 0.13 |
| *Dp(1;Y)BSC88* | (3C6-D2)-3E4 | 590 | 1834 | 541 | 7 | 190 | 0.09 |
| *Dp(1;Y)BSC90* | (3D5-3E4)-3E4 | 161 | 628 | 155 | 20 | 129 | 0.17 |
| *Dp(1;Y)BSC214* | (2E2-2F2)-2F6 | 120 | 984 | 252 | 12 | 199 | 0.17 |

\* Salivary gland chromosome bands. Left breakpoints are estimated between bands in parentheses.
\*\* Length of homology was calculated using the leftmost breakpoint.

### Direct observation of pairing between euchromatic homology on the X and Y

To directly ask if pairing was occurring between the euchromatic sequences on the *In(1)sc^4L^sc^8R^* and *Dp(1;Y)s*, we designed a FISH assay to cytologically visualize sex chromosome pairing in spermatocytes at early prophase I (S1-S2). Spermatocytes with diameters between 10 and 20 microns were selected because at this size they are considered to be in S1-S2a stage (Cenci *et al*. 1994) where pairing is observed (Vazquez, Belmont, and Sedat 2002). To assess pairing, a single copy X probe (Atto-594-Red) was hybridized to both the intact X and the X euchromatin duplicated on the *Dp(1;Y)*. Because both pairing and sister chromatid cohesion is lost as spermatocytes mature (Vazquez, Belmont, and Sedat 2002), a control chromosome 2 probe (Atto-488-Green) was used to assure the cells observed had not progressed beyond S2. Cells with two or more green signals were not scored as they may have already begun their progression to S3 when homologs no longer exhibit pairing. The X and Y were deemed paired when one red signal was present or two distinct signals were present that were less than 0.8 microns apart (Joyce *et al*. 2013).

There are two potential errors in this meiotic pairing assay that must be considered. First, there can be a slight asynchrony in the loss of pairing and sister chromatid cohesion on different chromosomes at the end of S2. Thus, some cells were predicted to be observed in which the X and Y had indeed paired, but sex chromosome pairing or sister chromatid cohesion had been lost prior to loss of pairing at the control autosomal site. This occurrence would have led to a false negative scoring of these cells as unpaired. To estimate how often this occurred, we hybridized the same probes to spermatocytes of males with *wildtype* sex chromosomes, so the red probe would only hybridize to the X. Ten percent of cells of the selected size in such males had one autosome signal and two X signals representing sister chromatid separation. This means that we may be underestimating pairing frequencies by as much as 10%.

Second, false positives in which pairing is erroneously scored are expected to occur by chance overlap of unpaired X signals. To estimate how often this occurs, we counted the number of spermatocytes that had overlap (within 0.8 µm) of the X and autosome signals in *Dp(1;Y)* males. Five percent of spermatocytes showed overlap of X and autosome signals. Overall, based on these two error rates, our measured frequencies may overestimate pairing by roughly five percent. Considering both sources of error, we expect that our overall estimates of pairing may be up to 5% less that the actual pairing frequencies.

Although *Dp(1;Y)*s varied in their ability to segregate from *In(1)sc^4L^sc^8R^*, all duplicated euchromatic sequences showed similar ability to pair with the homologous sequences on the intact X (Table 5). Considering our potential errors in estimation of pairing, some sequences showed nearly complete pairing. These results indicate that the observed differences in segregation of the various *Dp(1;Y)*s from the X could not be accounted for by differences in pairing ability (Table 5), but rather that pairing at some sites led to better segregation, possibly because of a greater ability to remain conjoined. To examine this possibility, we estimated that frequency at which paired chromosomes ultimately segregated properly for five different *Dp(1;Y)* genotypes. To avoid complications of meiotic drive, these estimates were based on direct measurements of pairing and segregation by FISH (see Materials and Methods). The abilities of the five *Dp(1;Y)*s to disjoin differed and showed the same trend with respect to the centromere proximity (Figure 6). These estimates supported our previous conclusion that the more proximal to the centromere the homology was on the X, the better its ability to direct segregation.

**Figure 6.**
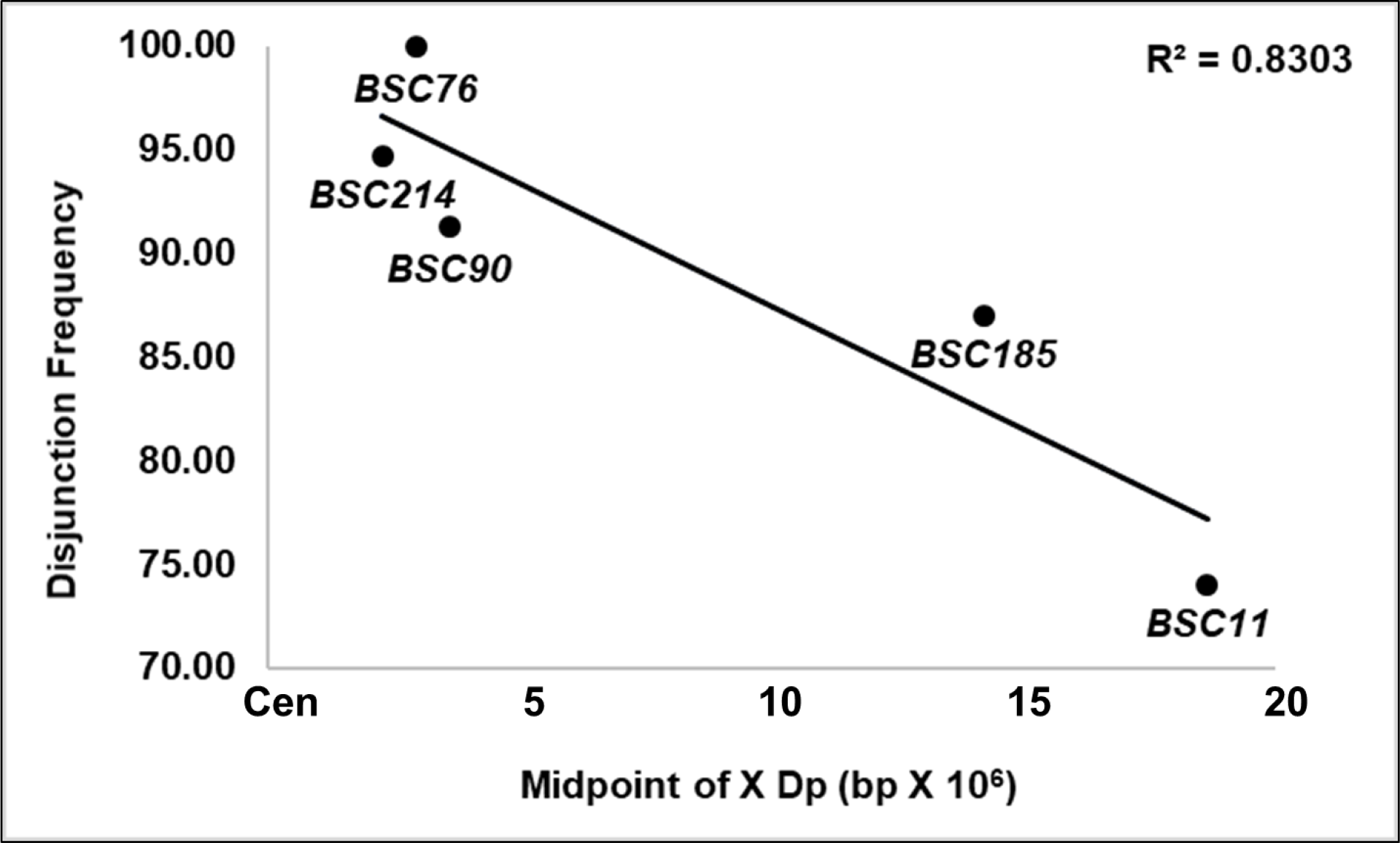
Frequency of disjunction of paired *In(1)sc^4L^sc^8R^* and *Dp(1;Y)BSC* chromosomes vs. genomic sequence position of the X homology.

**TABLE 5.** XY pairing in S1-S2 primary spermatocytes.

| X | Y | # Cells Scored | % Paired | % Paired that Disjoined * |
| --- | --- | --- | --- | --- |
| *wildtype* | *Dp(1;Y)BSC76* | 195 | 91.8 | ND |
| *In(1)sc<sup>4L</sup>sc<sup>8R</sup>* | *Dp(1;Y)BSC76* | 202 | 78.2 | 100.0 |
| *In(1)sc<sup>4L</sup>sc<sup>8R</sup>* | *Dp(1;Y)BSC214* | 236 | 93.2 | 94.7 |
| *In(1)sc<sup>4L</sup>sc<sup>8R</sup>* | *Dp(1;Y)BSC90* | 204 | 92.2 | 91.3 |
| *In(1)sc<sup>4L</sup>sc<sup>8R</sup>* | *Dp(1;Y)BSC185* | 215 | 73.5 | 87.0 |
| *In(1)sc<sup>4L</sup>sc<sup>8R</sup>* | *Dp(1;Y)BSC11* | 213 | 84.0 | 74.0 |

\* See Materials and Methods for calculation

We next asked if the ability of these euchromatic sequences to pair was dependent on the lack of the native X rDNA pairing sites. One possibility was that pairing might normally occur only at the rDNA if it had the ability to outcompete other homologies for limited pairing proteins. To test this, we measured pairing between an X chromosome bearing rDNA and *Dp(1;Y)BSC76* and found that pairing at the euchromatic homology was not diminished (Table 5). This shows pairing at the rDNA did not compete with pairing at the euchromatic homology.

### Effects of *tef* and *snm* on euchromatin-mediated sex chromosome segregation

We next used our pairing system to examine the requirements for the conjunction proteins Tef, MNM and SNM. Tef is normally required to maintain conjunction between autosomes, has no effects on sex chromosome segregation, and has proposed to be autosome-specific (Tomkiel, Wakimoto, and Briscoe 2001). However, because autosomal pairing sites are euchromatic and sex chromosome pairing sites are normally heterochromatic, the autosomal specificity of Tef may actually reflect a specificity for euchromatin. To test this possibility, we used the *In(1)sc^4L^sc^8R^ / Dp(1;Y)* pairing system to determine if Tef was required for euchromatic sex chromosome conjunction and segregation. First, we confirmed that the *In(1)sc^4L^sc^8R^* chromosome behavior was not altered in a *tef* background. We monitored sex chromosome NDJ of *In(1)sc^4L^sc^8R^ / y^+^ Y* males bearing a *tef* mutation and found that sex chromosome missegregation rates were statistically the same for *tef / + vs tef*, ((p > 0.95), Table 6). Next, we compared sex chromosome NDJ from *In(1)sc^4L^sc^8R^ / Dp(1;Y)BSC76* males homozygous or heterozygous for *tef*, and results did not differ statistically for *tef / + vs tef*, ((p > 0.50), Table 6).

**TABLE 6.**
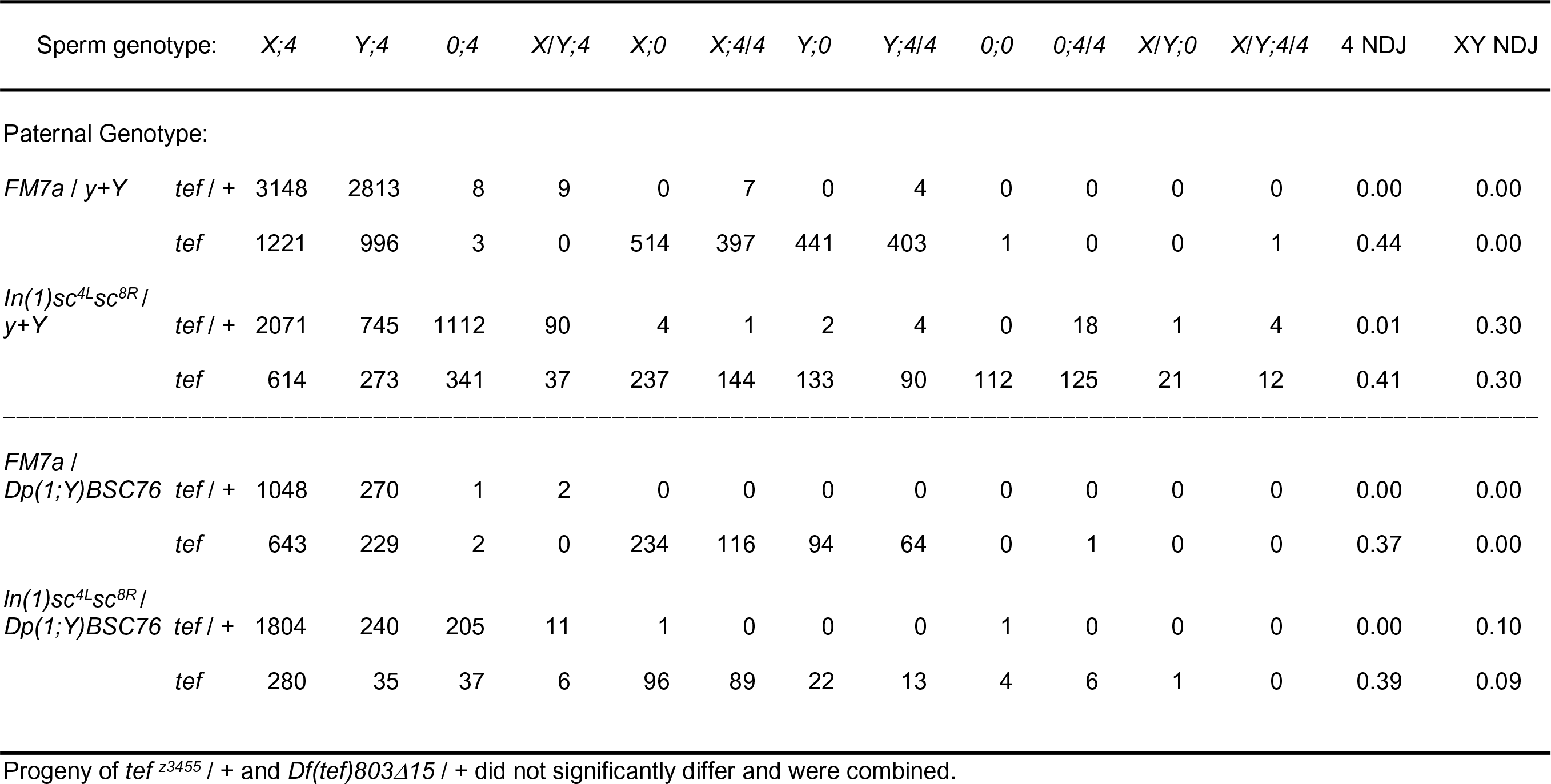
Effect of tef ^*z3455*^ / *Df(tef)803Δ15* on XY segregation in *In(1)sc^4L^sc^8R^* / *Dp(1;Y)BSC76* males

These results suggest that Tef is indeed autosome-specific and is not required for conjunction of these X euchromatic homologies.

We similarly attempted to test the requirements for MNM and SNM to establish conjunction between X euchromatic homologies. Unfortunately, we were unable to perform the same test. For unknown reasons, *In(1)sc^4L^sc^8R^ / Dp(1;Y)* males homozygous for *mnm* or *snm* were sterile. This was true for all alleles tested both as homozygotes and transheterozygotes (*snm^z0317^*, *snm^z2138^*, *mnm^z5578^*, *mnm^z3298^*, and *mnm^z3401^*). *X / Dp(1;Y); mnm* males were also sterile; however, we were able to assay NDJ in *X / Dp(1;Y); snm* males (i.e. males bearing a *wildtype* X). As SNM is necessary for conjunction at the rDNA, we reasoned that any segregation of the X from the *Dp(1;Y)* observed in *snm* males could be attributed to the behavior of the X euchromatic homologies. Therefore, we compared sex chromosome NDJ frequencies from *snm* or *snm / +* males bearing *Dp(1;Y)BSC76* or *Dp(1;Y)BSC67*.

Sex chromosome segregation in *X / Dp(1;Y)BSC76; snm* males was randomized, and not significantly different from control *X / B^s^ Y y^+^; snm* males (p > 0.75, Table 7). NDJ in *X / Dp(1;Y)BSC67; snm* males was actually slightly higher than in control *snm / +* males (p < 0.05). Whereas in previous crosses, *In(1)sc^4L^sc^8R^ / Dp(1;Y)BSC76* and *In(1)sc^4L^sc^8R^ / Dp(1;Y)BSC67* showed different NDJ frequencies, no differences were observed here (p > 0.25). These data indicate that SNM is required to mediate conjunction between X chromosome euchromatin.

**Table 7.** Effect of *snm^z0317^* / *snm^z2138^* on XY segregation in *X / Dp(1;Y)BSC* males.

| Sperm genotype: |  | X;4 | Y;4 | 0;4 | X/Y;4 | X;0 | Y;0 | 0;0 | X/Y;0 | 4 NDJ | 0/(X+0) |
| --- | --- | --- | --- | --- | --- | --- | --- | --- | --- | --- | --- |
| Parental Genotype: |  |  |  |  |  |  |  |  |  |  |  |
| $X / B^s Y y+$ | $snm^{z0317} / +$ | 1204 | 858 | 1 | 0 | 0 | 0 | 0 | 0 | 0.00 | 0.00 |
| | $snm$ | 425 | 274 | 386 | 182 | 112 | 104 | 121 | 98 | 0.26 | 0.49 |
| $X / Dp(1;Y)BSC76$ | $snm^{z0317} / +$ | 1296 | 385 | 3 | 0 | 0 | 0 | 0 | 0 | 0.00 | 0.00 |
| | $snm$ | 387 | 113 | 413 | 126 | 123 | 37 | 106 | 53 | 0.24 | 0.50 |
| $X / Dp(1;Y)BSC67$ | $snm^{z0317} / +$ | 881 | 217 | 20 | 7 | 0 | 0 | 0 | 0 | 0.00 | 0.02 |
| | $snm$ | 154 | 37 | 170 | 40 | 29 | 10 | 48 | 11 | 0.20 | 0.54 |

## DISCUSSION

The *Drosophila* male is an interesting model in which to study meiosis because homologs do not recombine and thus they lack the canonical mechanism of homolog attachment and segregation. It is also of particular interest because it was the first organism in which specific sequences were identified that function as meiotic pairing sites. A 240 bp sequence within the IGS of the rDNA is sufficient for pairing and segregation of the X from the Y (McKee and Karpen 1990; McKee, Lumsden, and Das 1993; McKee, Habera, and Vrana 1992). Although the X and the Y share significant sequence homology other than these IGS sequences in both the rDNA cistrons and at the stellate/crystal loci (Livak 1990), these homologies do not seem to promote pairing and segregation. Lack of pairing at other homologies suggested that there was a unique property of the IGS sequences with respect to sex chromosome meiotic pairing.

Similarly, there appeared to be some specificity to which autosomal sequences could function as “pairing sites”. Euchromatic segments of chromosome 2 translocated to the Y are capable of pairing and directing segregation from the intact chromosome 2 homolog, but a translocated segment of chromosome 2 heterochromatin is not (McKee, Lumsden, and Das 1993). Likewise, rearranged autosomal homologs that share only heterochromatic homologies do not pair and segregate from each other (Yamamoto 1979; Hilliker, Holm, and Appels 1982). These studies raised the question as to how the cell restricts pairing to specific sequences.

### Are there specific “pairing sites” in male meiosis?

Our work here suggests an alternative interpretation of these previous results. Prior observations of meiotic pairing were made during late prophase I, prometaphase I, and/or metaphase I (Yamamoto 1979; McKee, Habera, and Vrana 1992; McKee and Karpen 1990; McKee, Lumsden, and Das 1993). In these studies, chromosomes were judged as paired only if associations were observed in these later stages, and as such, failed to distinguish between the processes of pairing and conjunction.

Here, we have separately examined pairing and segregation (and by inference conjunction) utilizing a series of *Dp(1;Y)*s (Cook *et al*. 2010) and the rDNA-deficient *In(1)sc^4L^sc^8R^* X chromosome. Using *in situ* hybridization in combination with genetic tests of chromosome transmission, we were able to directly observe meiotic pairing independently of conjunction and assay its relationship to segregation. Our results indicate that 13 different Y chromosome rearrangements bearing X euchromatic homology are capable of pairing with the X. Rather than being limited to specific sequences, we suggest that pairing in males, as in other systems, may simply be homology-based. This possibility is consistent with observations that autosomal heterochromatic repeats are indeed paired in early prophase I (Tsai, Yan, and McKee 2011), and that *lacI* repeats inserted in 13 different euchromatic positions are all paired in early prophase I (Vazquez, Belmont, and Sedat 2002).

We found that all homologous segments tested paired with high fidelity (>74%). No relationship between homology length and pairing ability was observed, which means that either (1) pairing sites are not evenly distributed along the X chromosome or (2) the duplicated sequences tested (∼700 Kbp – 1500 Kbp) were all above the minimum threshold required for efficient pairing. We conclude that either all euchromatin can pair, or that pairing sites are distributed throughout the euchromatin.

To further address if there are minimal sequence requirements for XY pairing, we subdivided a duplicated euchromatic sequence into two smaller 120 Kb and 161 Kb fragments. We found that both sequences paired equally well, implying that the subdivided segment contains at least two sequences capable of pairing. Further analysis using deletions of these duplicated regions will be necessary to determine if pairing occurs at all euchromatin or if there are unique pairing sites within each tested region. In the absence of evidence for the latter, the most parsimonious explanation for our data is simply that all homologous sequences have the ability to pair.

### What determines conjunction in male meiosis?

If all homologous sequences can pair but not all remain associated and/or have the ability to direct segregation, then specific sequences may act as conjunction sites. Three proteins necessary for conjunction have been identified to date, MNM, SNM, and Tef. A putative MNM/SNM complex is required for conjunction for all bivalents, whereas Tef only affects conjunction between autosomal homologs (Thomas *et al*. 2005). By examining the pairing behavior of integrated *lacO* sites, it was concluded that mutants in *mnm* and *snm* do not disrupt pairing in S1 (Thomas *et al*. 2005), whereas the effects of *tef* mutants on pairing have not yet been examined. Both MNM and SNM localize to the 240 bp IGS repeats embedded within the rDNA cistrons (Thomas and McKee 2007). Tef is needed to localize MNM (and presumably SNM) to sites along the autosomes (Thomas *et al*. 2005). Whereas Tef binding sites have yet to be identified, the existence of three canonical C2H2 zinc fingers in Tef suggest that there may indeed be a consensus sequence for establishing conjunction on autosomes (Arya *et al*. 2006).

In our system, we examined the ability of X chromosome homologies to remain conjoined and thereby direct segregation. It was possible that these sequences lacked the MNM/SNM binding sites present in IGS sequences and also the autosomal binding sites potentially recognized by Tef. We wondered which, if any, of these proteins might be involved in mediating conjunction. We first tested if *tef* mutations had any effect on X / *Dp(1;Y)* segregation. Although *tef* mutations show an autosome-specificity, it was possible that this specificity reflected a euchromatin-specific function that did not affect the normally heterochromatic XY conjunction. If this were the case, we might have expected *tef* mutations to disrupt the euchromatin-mediated XY conjunction. We found, however that Tef was not required suggesting that Tef is indeed specific for autosomes.

We next sought to test the requirements for MNM and SNM. While SNM and MNM show binding specificity to IGS sequences (Thomas and McKee 2007), the exact binding sites within the IGS have not been determined. It is not known if potential binding sequences might also be distributed throughout X euchromatin.

Unfortunately, we were unable to test the role of MNM because for an unknown reason, MNM mutants in combination with the sex chromosome rearrangements were sterile. However, we were able to test SNM, and indeed, found it to be required for segregation in our X-Y euchromatic pairing system. This result shows that SNM is necessary for conjunction between X euchromatin and suggests that sequences sufficient for SNM binding are present in X euchromatin. Because Tef is not required, the mechanism of SNM binding to the X euchromatin likely differs from the mechanism by which SNM binds to the autosomes. There may be homology to IGS sequences in the X euchromatin that directly bind SNM, although we could not identify extensive homology using BLAST (Altschul *et al*. 1990). Interestingly, there is a cluster of IGS-like sequences present on chromosome 3R that share almost 90% identity to the rDNA IGS repeats (FLYBASE). Polymorphisms that differentiate these sequences from the X rDNA IGS sequences may be critical in determining SNM binding.

An alternative explanation for SNM-mediated conjunction at X euchromatin is that *In(1)sc^4L^sc^8R^* may have a small number of remaining IGS sequences. One or two IGS sequences on their own may not be sufficient for establishing pairing but may be sufficient for mediating conjunction if pairing via euchromatin occurred *in cis*.

### Centromere-proximal sequences are more effective at directing segregation

Interestingly, although we found all homologous sequences paired with similar fidelity, not all sequences behaved the same in the ability to direct segregation. Pairings between centromere proximal sequences were better at directing homolog segregation. A similar observation was made for the segregation of *Dp(2;Y)*s from intact chromosome 2 homologs. Euchromatic homology found to be most effective at directing segregation was the histone locus, which resides on 2R adjacent to the centromere (McKee, Lumsden, and Das 1993).

Why might centromere-proximal association demonstrate a greater frequency of proper segregation? One possibility is that pairing close to the remaining heterochromatin of the *In(1)sc^4L^sc^8R^* X may be more effective at establishing conjunction at cryptic IGS sequences. Proximal pairing may be better at bringing such sites on homologs close enough to facilitate conjunction. Alternatively, centromere-proximal attachments could simply be better at establishing tension across the bivalent at metaphase I. Tension is important for stabilizing kinetochore attachments necessary for establishing bipolar orientation (Salmon and Bloom 2017). In many systems, when tension is not present at kinetochores because of insufficient microtubule attachment, a metaphase arrest is triggered (Nicklas *et al*. 2001). In male *Drosophila*, however, activation of this checkpoint by unpaired chromosomes merely delays the transition to anaphase I (Rebollo and Gonzalez 2000). It is conceivable that meiosis would proceed through anaphase I even if the XY bivalent had not formed stable bipolar attachments, leading to NDJ. This possibility may explain why the centromere-proximal rDNA locus evolved as the native XY pairing site.

In summary, our examination of XY euchromatic pairing suggests some fundamental differences in the previous models of meiotic pairing and conjunction in male flies. Rather than pairing being limited to specific sequences, we propose that the simplest model is that all homologous sequences can pair, and only a subset of homologies function as conjunction sites during meiosis I. The repeats with the IGS sequences of the rDNA are most likely conjunction sites which serve to bind the conjunction proteins MNM and SNM (Thomas and McKee 2007), and a putative complex of these proteins with Tef may localize to conjunction sites within autosomal euchromatin. Conjunction sites may be able to pair, but not all pairing sites may be capable of establishing conjunction.

Our assay promises to be useful to further define requirements for meiotic pairing. Deletion analysis of euchromatic region may delimit the minimal sequences required for pairing and determine whether specific sequences are required for pairing and/or conjunction.

## ACKNOWLEDGEMENTS & FUNDING

We thank Dr. Barbara Wakimoto for providing fly lines. This work was supported by National Institutes of Health grant R15GM119055 to J.E.T and the University of North Carolina at Greensboro, UNCG Graduate Student Research Award to C.A.H., and UNCG Undergraduate Research Awards to K.H., and A.B.

## REFERENCES

1. Altschul, S. F., W. Gish, W. Miller, E. W. Myers, and D. J. Lipman. 1990. ‘Basic local alignment search tool’, J Mol Biol, 215: 403–10.

2. Arya, G. H., M. J. Lodico, O. I. Ahmad, R. Amin, and J. E. Tomkiel. 2006. ‘Molecular characterization of teflon, a gene required for meiotic autosome segregation in male Drosophila melanogaster’, Genetics, 174: 125–34.

3. Bahler, J., T. Wyler, J. Loidl, and J. Kohli. 1993. ‘Unusual nuclear structures in meiotic prophase of fission yeast: a cytological analysis’, J Cell Biol, 121: 241–56.

4. Beliveau, B. J., N. Apostolopoulos, and C. T. Wu. 2014. ‘Visualizing genomes with Oligopaint FISH probes’, Curr Protoc Mol Biol, 105: Unit 14 23.

5. Cenci, G., S. Bonaccorsi, C. Pisano, F. Verni, and M. Gatti. 1994. ‘Chromatin and microtubule organization during premeiotic, meiotic and early postmeiotic stages of Drosophila melanogaster spermatogenesis’, Journal of Cell Science, 107: 3521–34.

6. Cook, R. K., M. E. Deal, J. A. Deal, R. D. Garton, C. A. Brown, M. E. Ward, R. S. Andrade, E. P. Spana, T. C. Kaufman, and K. R. Cook. 2010. ‘A new resource for characterizing X-linked genes in Drosophila melanogaster: systematic coverage and subdivision of the X chromosome with nested, Y-linked duplications’, Genetics, 186: 1095–109.

7. Cooper, K. W. 1959. ‘Cytogenic analysis of major heterochromatic elements (especially Xh and Y) in *Drosophila melanogaster*, and the theory of ‘heterochromatin’’, Chromosoma, 10: 535–88.

8. Giroux, C. N., M. E. Dresser, and H. F. Tiano. 1989. ‘Genetic control of chromosome synapsis in yeast meiosis’, Genome, 31: 88–94.

9. Gramates, L. S., S. J. Marygold, G. D. Santos, J. M. Urbano, G. Antonazzo, B. B. Matthews, A. J. Rey, C. J. Tabone, M. A. Crosby, D. B. Emmert, K. Falls, J. L. Goodman, Y. Hu, L. Ponting, A. J. Schroeder, V. B. Strelets, J. Thurmond, P. Zhou, and Consortium the FlyBase. 2017. ‘FlyBase at 25: looking to the future’, Nucleic Acids Res, 45: D663–D71.

10. Hilliker, A. J., D. G. Holm, and R. Appels. 1982. ‘The relationship between heterochromatic homology and meiotic segregation of compound second autosomes during spermatogenesis in Drosophila melanogaster’, Genet Res, 39: 157–68.

11. Joyce, E. F., N. Apostolopoulos, B. J. Beliveau, and C. T. Wu. 2013. ‘Germline progenitors escape the widespread phenomenon of homolog pairing during Drosophila development’, PLoS Genet, 9: e1004013.

12. Kemp, B., R. M. Boumil, M. N. Stewart, and D. S. Dawson. 2004. ‘A role for centromere pairing in meiotic chromosome segregation’, Genes Dev, 18: 1946–51.

13. Livak, K. J. 1990. ‘Detailed structure of the Drosophila melanogaster stellate genes and their transcripts’, Genetics, 124: 303–16.

14. Lohe, A. R., and D. L. Brutlag. 1987. ‘Adjacent satellite DNA segments in Drosophila structure of junctions’, J Mol Biol, 194: 171–9.

15. MacQueen, A. J., C. M. Phillips, N. Bhalla, P. Weiser, A. M. Villeneuve, and A. F. Dernburg. 2005. ‘Chromosome sites play dual roles to establish homologous synapsis during meiosis in C. elegans’, Cell, 123: 1037–50.

16. McKee, B. 1984. ‘Sex Chromosome Meiotic Drive in DROSOPHILA MELANOGASTER Males’, Genetics, 106: 403–22.

17. McKee, B. D., L. Habera, and J. A. Vrana. 1992. ‘Evidence that intergenic spacer repeats of Drosophila melanogaster rRNA genes function as X-Y pairing sites in male meiosis, and a general model for achiasmatic pairing’, Genetics, 132: 529–44.

18. McKee, B. D., and G. H. Karpen. 1990. ‘Drosophila ribosomal RNA genes function as an X-Y pairing site during male meiosis’, Cell, 61: 61–72.

19. McKee, B. D., S. E. Lumsden, and S. Das. 1993. ‘The distribution of male meiotic pairing sites on chromosome 2 of Drosophila melanogaster: meiotic pairing and segregation of 2-Y transpositions’, Chromosoma., 102: 180–94.

20. McKim, K. S., B. L. Green-Marroquin, J. J. Sekelsky, G. Chin, C. Steinberg, R. Khodosh, and R. S. Hawley. 1998. ‘Meiotic synapsis in the absence of recombination’, Science, 279: 876–8.

21. Nicklas, R. B., J. C. Waters, E. D. Salmon, and S. C. Ward. 2001. ‘Checkpoint signals in grasshopper meiosis are sensitive to microtubule attachment, but tension is still essential’, J Cell Sci, 114: 4173–83.

22. Peacock, W. J., G. L. Miklos, and D. J. Goodchild. 1975. ‘Sex chromosome meiotic drive systems in Drosophila melanogaster I. Abnormal spermatid development in males with a heterochromatin-deficient X chromosome (sc-4sc-8)’, Genetics, 79: 613–34.

23. Rebollo, E., and C. Gonzalez. 2000. ‘Visualizing the spindle checkpoint in Drosophila spermatocytes’, EMBO Rep, 1: 65–70.

24. Ren, X., L. Eisenhour, C. Hong, Y. Lee, and B. D. McKee. 1997. ‘Roles of rDNA spacer and transcription unit-sequences in X-Y meiotic chromosome pairing in Drosophila melanogaster males’, Chromosoma, 106: 29–36.

25. Ritossa, F. M. 1968. ‘Unstable redundancy of genes for ribosomal RNA’, Proc Natl Acad Sci U S A, 60: 509–16.

26. Ritossa, F. M.. 1976. ’The Bobbed Locus.’ in M. Ashburner and A. Novitski (eds.), The Genetics and Biology of Drosophila (Academic PRess: London).

27. Salmon, E. D., and K. Bloom. 2017. ‘Tension sensors reveal how the kinetochore shares its load’, Bioessays, 39.

28. Sato, A., B. Isaac, C. M. Phillips, R. Rillo, P. M. Carlton, D. J. Wynne, R. A. Kasad, and A. F. Dernburg. 2009. ‘Cytoskeletal forces span the nuclear envelope to coordinate meiotic chromosome pairing and synapsis’, Cell, 139: 907–19.

29. Sun, M. S., J. Weber, A. C. Blattner, S. Chaurasia, and C. F. Lehner. 2019. ‘MNM and SNM maintain but do not establish achiasmate homolog conjunction during Drosophila male meiosis’, PLoS Genet, 15: e1008162.

30. Tartof, K. D. 1974. ‘Unequal mitotic sister chromatin exchange as the mechanism of ribosomal RNA gene magnification’, Proc Natl Acad Sci U S A, 71: 1272–6.

31. Thomas, S. E., and B. D. McKee. 2007. ‘Meiotic pairing and disjunction of mini-X chromosomes in drosophila is mediated by 240-bp rDNA repeats and the homolog conjunction proteins SNM and MNM’, Genetics, 177: 785–99.

32. Thomas, S. E., M. Soltani-Bejnood, P. Roth, R. Dorn, J. M. Logsdon, Jr., and B. D. McKee. 2005. ‘Identification of two proteins required for conjunction and regular segregation of achiasmate homologs in Drosophila male meiosis’, Cell, 123: 555–68.

33. Tomkiel, J. E., B. T. Wakimoto, and A. Briscoe, Jr. 2001. ‘The teflon gene is required for maintenance of autosomal homolog pairing at meiosis I in male Drosophila melanogaster’, Genetics, 157: 273–81.

34. Tsai, J. H., R. Yan, and B. D. McKee. 2011. ‘Homolog pairing and sister chromatid cohesion in heterochromatin in Drosophila male meiosis I’, Chromosoma, 120: 335–51.

35. Vazquez, J., A. S. Belmont, and J. W. Sedat. 2002. ‘The dynamics of homologous chromosome pairing during male Drosophila meiosis’, Curr Biol, 12: 1473–83.

36. Wakimoto, B. T., D. L. Lindsley, and C. Herrera. 2004. ‘Toward a comprehensive genetic analysis of male fertility in Drosophila melanogaster’, Genetics, 167: 207–16.

37. Weiner, B. M., and N. Kleckner. 1994. ‘Chromosome pairing via multiple interstitial interactions before and during meiosis in yeast’, Cell, 77: 977–91.

38. Wynne, D. J., O. Rog, P. M. Carlton, and A. F. Dernburg. 2012. ‘Dynein-dependent processive chromosome motions promote homologous pairing in C. elegans meiosis’, J Cell Biol, 196: 47–64.

39. Yamamoto, M. 1979. ‘Cytological studies of heterochromatin function in the Drosophila melanogaster male: autosomal meiotic paring’, Chromosoma, 72: 293–328.

